# Dimensionality reduction by UMAP to visualize physical and genetic interactions

**DOI:** 10.1101/681726

**Authors:** Michael W. Dorrity, Lauren M. Saunders, Christine Queitsch, Stanley Fields, Cole Trapnell

**Author notes:** these authors contributed equally (Michael W. Dorrity, Lauren M. Saunders).

## Abstract

Dimensionality reduction is often used to visualize complex expression profiling data. Here, we use the Uniform Manifold Approximation and Projection (UMAP) method on published transcript profiles of 1484 single gene deletions of *Saccharomyces cerevisiae*. Proximity in low-dimensional UMAP space identifies clusters of genes that correspond to protein complexes and pathways, and finds novel protein interactions even within well-characterized complexes. This approach is more sensitive than previous methods and should be broadly useful as additional transcriptome datasets become available for other organisms.

---

A central goal of biological studies is the identification and characterization of proteins that act in a common cellular pathway. Efforts towards this goal have been greatly aided by large-scale perturbation analyses coupled with whole-transcriptome profiling, in which each gene’s transcriptional response to a perturbation is measured. If a sufficient database of expression profiles exists, then a pathway affected by an uncharacterized perturbation – such as a gene mutation, drug treatment or growth condition – can be described by matching the resultant profile to a known profile.^1^ For the yeast *Saccharomyces cerevisiae*, the expression profiles of a large number of individual yeast deletion mutants have been established and used to infer protein complexes and networks.^2–4^ Maximizing the utility of expression profiling approaches for inference of physical and genetic interactions requires ever larger such datasets. However, standard techniques, such as pairwise correlation, do not fully capture the variation available to link gene function as more dimensions are added from larger scale experiments. Therefore, techniques that reduce dimensionality of the data while maintaining relationships between genes are imperative for the inference of physical and genetic interactions in very large gene expression datasets.

Dimensionality reduction methods capture variability in a limited number of random variables to facilitate 2- or 3D-visualization of datasets with tens to thousands of dimensions. This approach is recognizable in the commonly used method of principal component analysis (PCA), which uses linear combinations of variables to generate orthogonal axes that efficiently capture the variation present in the data with fewer variables. Another approach, t-Distributed Stochastic Neighbor Embedding (t-SNE), carries out dimensionality reduction by analyzing similarity of points using a Gaussian distance in high dimensional space and projecting these data into a low dimensional space.^5^ A more recent method, uniform manifold approximation (UMAP), estimates a topology of the high dimensional data and uses this information to construct a low-dimensional representation that preserves relationships present in the data.^6^ UMAP has been particularly useful to precisely define cell types in mixed populations based on data from single-cell RNA-seq experiments^7–13^; it also performs well on other gold-standard datasets.^6,14^ Because UMAP is better able to preserve elements of the data structure from high dimensional space than similar outputs from t-SNE, it captures local relationships within distinct clusters in addition to global relationships between clusters.^14^ This feature is especially useful in the inference of gene relationships, which can be due to physical interaction, overlapping gene function, or coordinated contributions to a larger cellular process. Here, we show that the use of dimensionality reduction by UMAP on bulk expression profiling data of 1484 single gene mutants of *S. cerevisiae* links gene function in clusters at increasingly finer scales, corresponding to broad cellular activities, pathways, protein complexes and individual protein-protein interactions.

We assigned groups, or “clusters,” to deletion mutants with similar transcriptional responses using the Louvain community detection algorithm in low-dimensional UMAP space.^9^ While many single-cell transcriptomic studies use expression values from genes with the highest dispersion across individual cells, we took advantage of the completeness of bulk microarray data generated by Kemmeren et al.^3^ and used expression values for all 6170 genes to make a UMAP projection for subsequent clustering. This approach resolved 50 main clusters, with the number of deletion backgrounds assigned to each cluster ranging from 4 to 298 (median of 11). Clusters with >25 genes were subsequently re-clustered using similar parameters to define sub-clusters. The final dataset contains 171 clusters with a median of 8 genes per cluster.

A total of 194 characterized yeast complexes have at least two of their corresponding genes in the dataset of single deletions. For 40% of these complexes (78/194), we could assign two or more genes to the same cluster (examples of complexes in the initial set of 50 clusters in Figure 1A, additional complexes were separated in the re-clustered set (Figure 1B)). For example, the re-clustering of the original cluster 2, which is characterized by cell cycle and chromosome organization genes, resulted in more distinctly separating the Isw2-Itc1 chromatin remodeling complex, the Csm3-Tof1 S-phase checkpoint complex and the Oca S-phase histone activation complex (Figure 1B). Within this re-clustered set, multiple complexes could be found among genes within a single cluster, suggesting that these complexes may cooperatively contribute to chromosome cohesion and recombination (Figure 1B).

**Figure 1.**
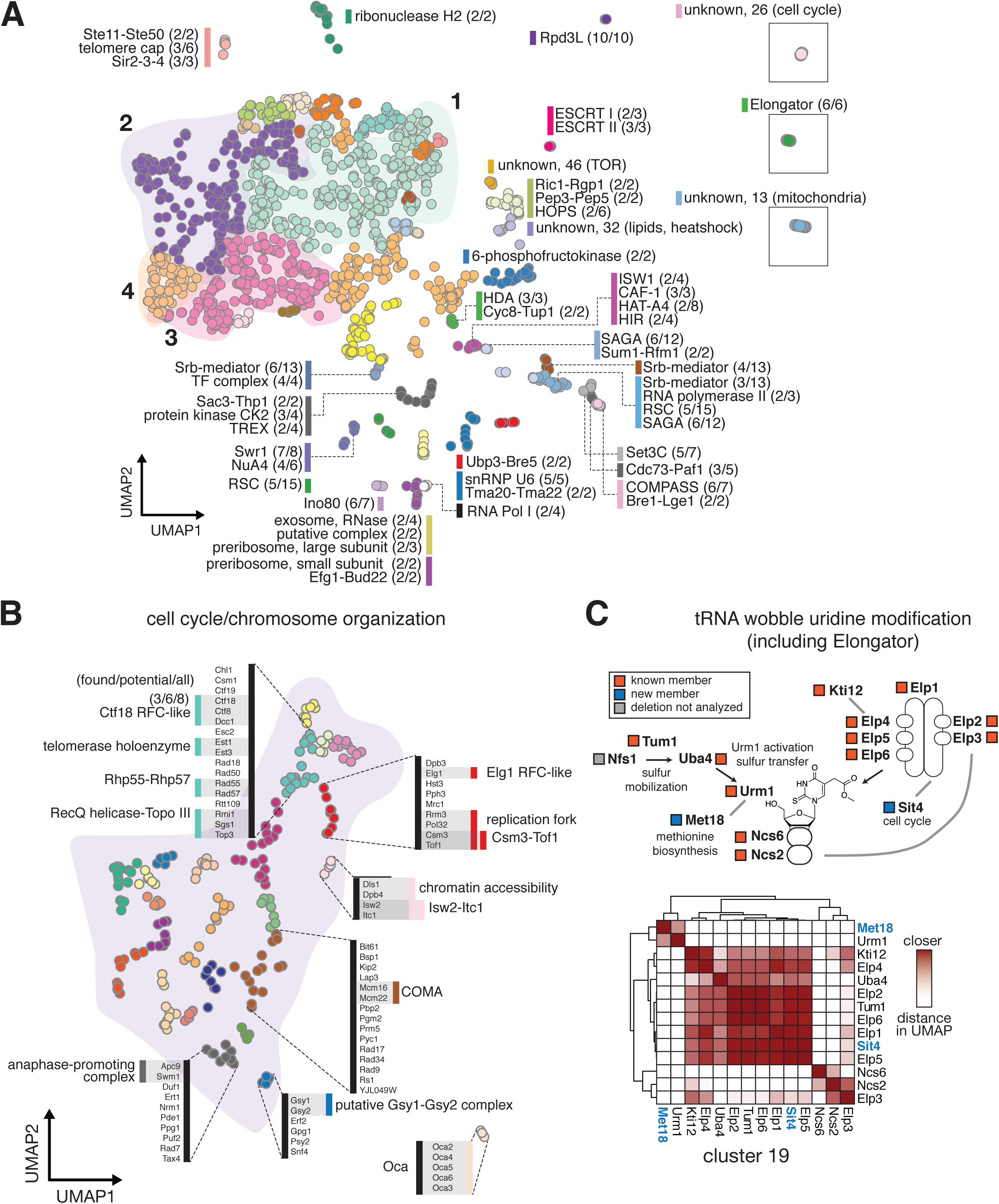
UMAP clusters single gene deletion transcriptomes according to shared function. (A) UMAP coordinates of 1484 single gene deletion strains clustered by similarity in transcriptional effects. The initial 50 individual clusters are each shown in a different color. Strains that comprise protein complexes are indicated alongside a bar colored according to cluster identity. Each complex is represented as a fraction: the number of complex members found in the cluster over the number of complex members in the set of 1484 mutants. Clusters with coordinates far from the main group are shown in boxes. Clusters without a known complex are marked as “unknown,” along with an arbitrary cluster number; these clusters are annotated with a broad GO term enriched in that cluster. (B) Cluster 2 shows more distinct groupings when re-clustered separately. Annotations as in (A). Cluster 2 as a whole was enriched for cell cycle and chromosome organization, with individual clusters corresponding to parts of this process. (C) The tRNA wobble uridine pathway, captured entirely within the cluster containing the Elongator complex (boxed green cluster in (A)). Complex members within this cluster are annotated with orange boxes, while new members are annotated in blue. One pathway member, Nfs1, was not present in the single gene deletion dataset. The heatmap represents fine-scale distances between each pair of points within the cluster. Darker shades of red indicate points nearer in UMAP space; hierarchical clustering was applied on this distance metric to group proteins within this pathway. Heterodimeric interactions, such as Ncs6-Ncs2 (bottom-right corner of heatmap), are nearer to each other than other members of the pathway. Novel members of this pathway (blue text) are grouped with other members based on their similarity of UMAP distance, and these new interactions are indicated with gray lines in the pathway diagram.

In some cases, members of individual complexes were assigned to separate clusters, suggesting sub-functionalization of components. For example, the 13-member mediator complex was found in three clusters (numbers 16, 34 and 41) containing 3, 6, and 4 members of mediator, respectively (Figure 1A). Cluster 16 (with 3 members) also contains members of SAGA and SWI/SNF complexes, and loss of mediator subunits in this cluster alters the transcription of amino acid metabolism genes and glucose transmembrane transporters (Supplemental Table 1); cluster 34 (with 6 members) contains galactose-responsive subunits of mediator; and cluster 41 (with 4 members) contains transcriptional-initiation-related mediator subunits. Here, UMAP preserves global relationships between clusters in addition to resolving proximal cluster members. For example, most chromatin remodeling complexes grouped in UMAP space, despite being present in separate clusters and containing unique local topologies (Figure 1A).

UMAP clustering identified the components of the pathway for tRNA wobble uridine modification and revealed two additional members that are likely to link metabolism and cell cycle to this process. One of these, Met18, has a human ortholog (MMS19) that functions in maturation of Fe-S cluster-containing proteins; the conserved yeast and human Elongator component Elp3 is one of these Fe-S proteins.^15^ The other new member, the PP2A phosphatase Sit4, is implicated in dephosphorylation of Elongator; its absence leads to tRNA modification defects.^16^

To assess whether UMAP distance captured known interactions as well as pairwise correlation, we used a dataset of 1060 protein interactions determined from co-immunoprecipitation followed by mass spectrometry.^2^ The UMAP clustering data captured these complexes more sensitively and with more precision than previous pairwise correlation-based metrics (AUC pairwise correlation = 0.73, AUC UMAP = 0.84, Figure 2A). UMAP distance also captured known interacting pairs better than distance in high-dimensional space (AUC = 0.56) and distance in PCA space (AUC = 0.70), suggesting that the UMAP dimensionality reduction itself adds value in the identification of interactions (Figure 2A, Supplemental Figure 1A).

**Figure 2.**
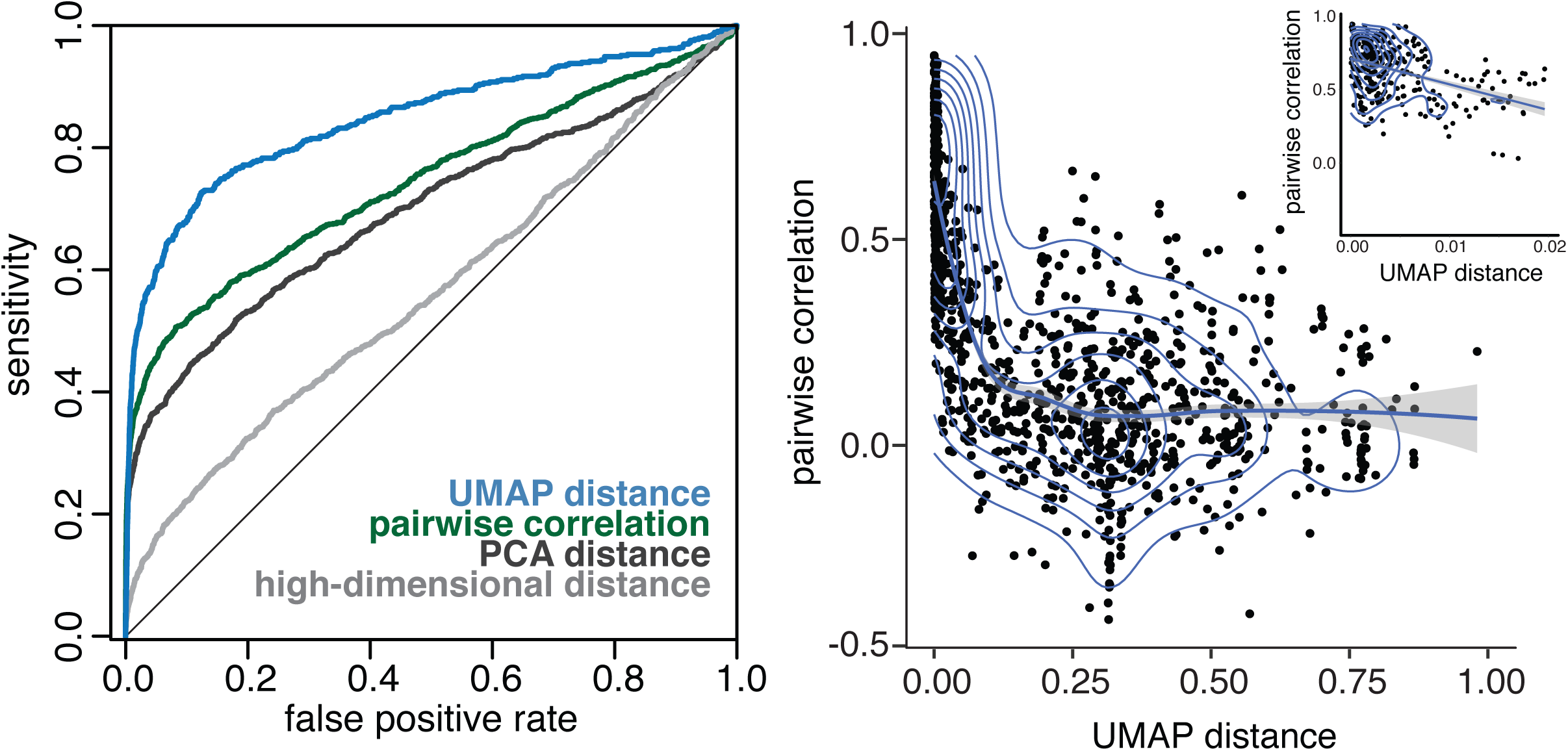
UMAP distance identifies protein-protein interactions more effectively than previous methods. (A) A receiver-operator curve showing the ability of UMAP distance to capture known protein-protein interactions (sensitivity) as a function of its false positive detection. UMAP distance (blue) performs better than pairwise correlation (green), PCA distance (dark grey), and high-dimensional distance (light grey) in identifying interactions. (B) For each protein-protein interaction, the distance between points in UMAP space was plotted against the pairwise correlation of that pair of transcriptomes. The density of points is indicated with blue lines. Inset in the upper right shows a zoomed-in portion of the x-axis; points with UMAP distance in this range are highly enriched for true interactions that are not captured by pairwise correlation.

Performing clustering in UMAP space ought to produce clusters containing more true interactions than distance in other spaces. To test whether similar results were obtained without UMAP dimensionality reduction, we clustered the data in PCA space. Clustering in PCA space identified 8/50 clusters with perfect overlap to UMAP clusters, and 34/50 overlapping by at least 50% (Supplemental Figure 1B).

To compare pairwise correlation with the UMAP approach, we calculated for each known interacting pair (1) the Pearson correlation of their deletion transcriptomes; and (2) the distance of those two genes in the UMAP space generated from using all deletion transcriptomes. Among these interacting pairs, UMAP distance and pairwise correlation are negatively correlated (Figure 2B). However, the increased sensitivity of UMAP distance to detect known interactions suggests that the discrepancies between UMAP distance and pairwise correlation might represent interactions that were previously overlooked. Based on a UMAP distance cutoff corresponding to a 5% FDR of known complex members (Inset - Figure 2B), we were able to identify 176 putative interactions that would not have been confidently called by previous approaches using pairwise correlations (PCC < 0.5); these interactions contain 86 unique genes, of which 77 show co-IP or yeast two-hybrid evidence for membership among 31 protein complexes, while the remaining 9 genes had no such evidence.

Since proximity in UMAP space tends to capture known interactions and shared function, distance in UMAP space could serve as a useful tool to investigate evolutionary questions about gene divergence. We calculated UMAP distance between 151 paralogous gene pairs in yeast and used this distance to characterize the functional divergence between each pair (Supplemental Figure 2A). Proximity of paralog pairs in UMAP space did not correspond to previous estimations of paralog divergence (Supplemental Figure 2B-C) based on synthetic genetic interaction (R=0.018) or Gene Ontology relationships (R =0.035).^17^ When paralogs show a negative genetic interaction – that is, deletion of both genes leads to lower fitness than expected – it is assumed that the two genes retain redundant functions. However, in 11 paralog pairs whose negative genetic interactions suggested redundant function, we showed distinct downstream effects on gene expression when each gene was deleted (Supplemental Figure 2B, 2D); these genes may have distinct effects on fitness in different environments.^18^ In these cases, a gene may retain the capacity to complement the essential function of its paralogous partner, while diverging sufficiently in function as revealed by the UMAP-based transcriptome analysis.

Despite successful clustering of many protein complexes and pathways of yeast, the UMAP approached nevertheless identified several clusters that did not obviously correspond to a complex or pathway. We used GO enrichment of differentially expressed genes in these clusters to interrogate their function: cluster 26 showed enriched terms for cell cycle, non-membrane-bound organelles, and prions; cluster 13 showed enrichment for mitochondrial function; cluster 46 showed enrichment for TOR signaling and aerobic respiration; cluster 32 showed enrichment for protein folding; and cluster 11 showed enrichment for heme binding. Differential expression analysis produced significant gene sets for all main and sub-clusters (Supplemental Table 1).

Because of its greater sensitivity than other approaches, as well as its ability to capture both local and global relationships, UMAP-based association of gene function adds value in the identification of protein complexes, pathways, and novel interactions in transcriptomic datasets. However, the utility of this method is dependent on the availability of high-quality profiling data from large-scale environmental or genetic perturbation experiments. As more datasets of this type become available, we expect that this approach, or similar dimensionality reduction techniques, will becoming increasingly useful in mapping protein complexes and pathways both within and across other species. The recent appearance of single-cell expression profiling data paired with CRISPR-induced mutations will be an especially useful source of data of this type, as these experiments include increasingly larger numbers of mutations.^19^ While many of the most useful applications of dimensionality reduction tend to arise from single-cell genomics, for which typical datasets necessitate approaches like UMAP to define relationships between cells, these approaches may also prove useful in visualizing the spatial relationships of biomolecules in tissues,^20^ genetic interactions, or relationships between human populations.^21^

## Methods

### Yeast single gene deletion transcriptome data

Growth-rate adjusted microarray expression values derived from limma modeling by Kemmeren et al.^2^ were used as input data. All 1484 single-gene deletion strains from this dataset were used for subsequent dimensionality reduction.

### UMAP dimensionality reduction and clustering

We used Monocle3 (v2.99.3) to perform Uniform Manifold Approximation and Projection (UMAP)^6^ to project single-gene deletion strains into two dimensions and performed Louvain clustering^22^ using default parameters (except, reduceDimension: reduction_method=UMAP, metric=cosine, n_neighbors=10, min_dist = 0.05; clusterCells: method=louvain, res=1e-4, k=3). Expression values from all 6170 yeast genes were given as input to Principal Component Analysis (PCA). The top 100 principal components were then used as input to UMAP for generating 2D projections of the data. For subclustering, main clusters 1 – 10 were each individually processed using top 25 principal components in the subset data as input to UMAP dimensionality reduction and Louvain clustering.

### Differentially expressed genes per cluster

Gene expression values for single-gene deletions within a cluster were compared to the background set of all deletions. Differentially expressed genes for each cluster were calculated using the differentialGeneTest() function in Monocle. Because the expression datasets were microarray-derived rather than count-based RNA-seq data, the ‘gaussianff’ expression family was used; significance values were corrected for genomic inflation factors using lamba gc (test.stat = qchisq(1.0 - p.val, df = 1)’’lambda.gc = median(dea_df$test.stat) / qchisq(0.5, 1).^23^

### Benchmarking with known interacting pairs

To test the ability of UMAP distance, and other distance metrics, to capture known interactions, we used a curated consensus set of protein complexes derived from two large, high-throughput mass spectrometry datasets and GO interactions.^2^ The consensus set was transformed into a pairwise Boolean interaction matrix based on whether or not each pair had been observed together in the known complex set. Using the subset of pairs that were found in the set of 1484 single gene deletion transcriptome datasets, for each gene pair, we calculated Euclidean distance in UMAP space, and used cumulative distance cutoffs to quantify the number of true and false interacting pairs and generate a receiver operating characteristic (ROC) curve.

### Code availability

All input data and scripts used for dimensionality reduction and clustering are available through github (https://github.com/cole-trapnell-lab/yeast_umap).

## Acknowledgements

We thank J. Packer for advice on differential expression analysis.

---

**Supplemental Figure 1.**
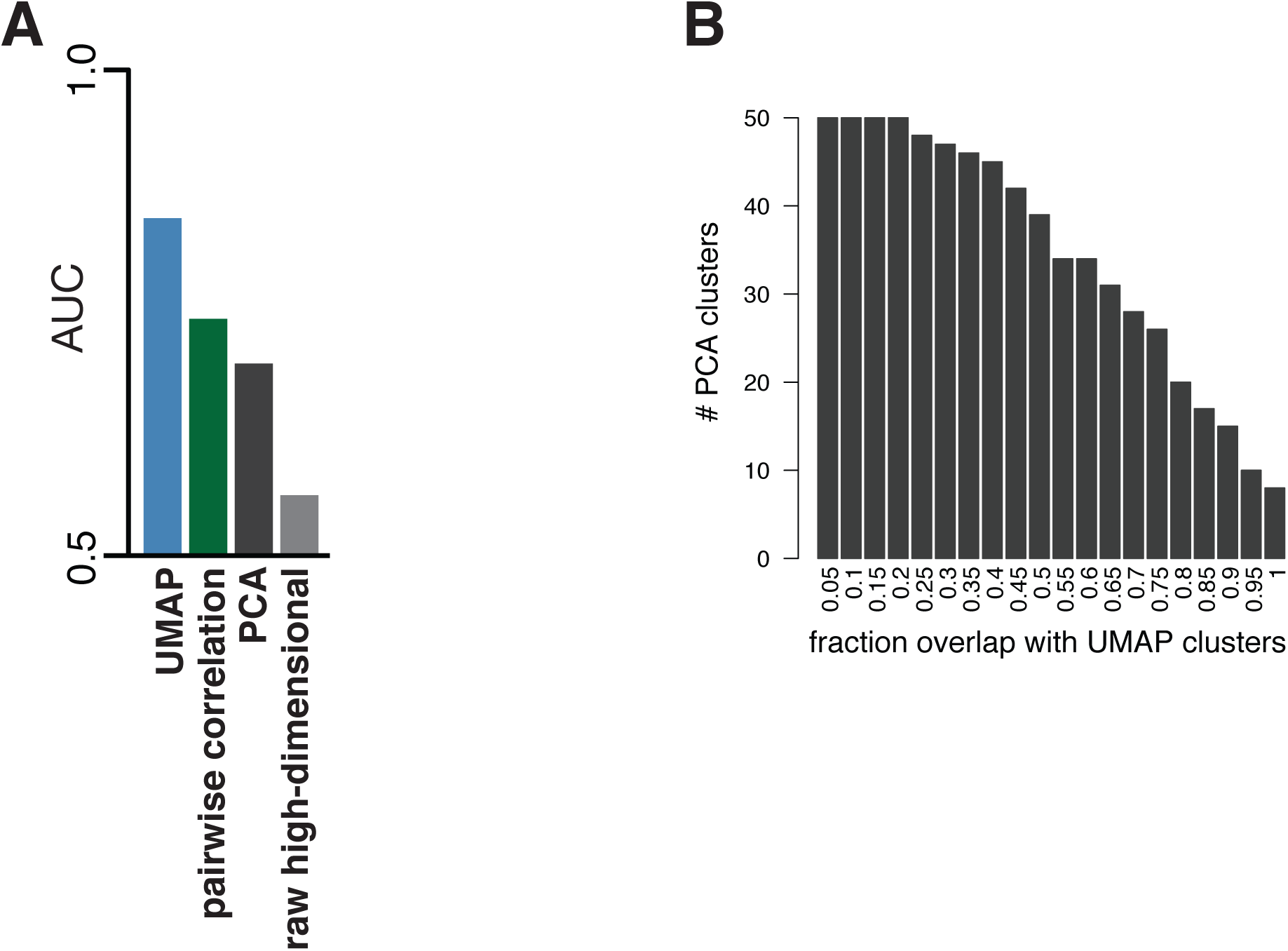
UMAP adds value in identification of true interactions compared to other methods. (A) Values for area under the curves (AUC) for ROC analysis in Figure 2A. UMAP substantially outperforms other metrics in the identification of true protein-protein interactions. (B) Clustering to the same total number of clusters in PCA space returns few clusters strongly overlapping the clusters in UMAP space.

**Supplemental Figure 2.**
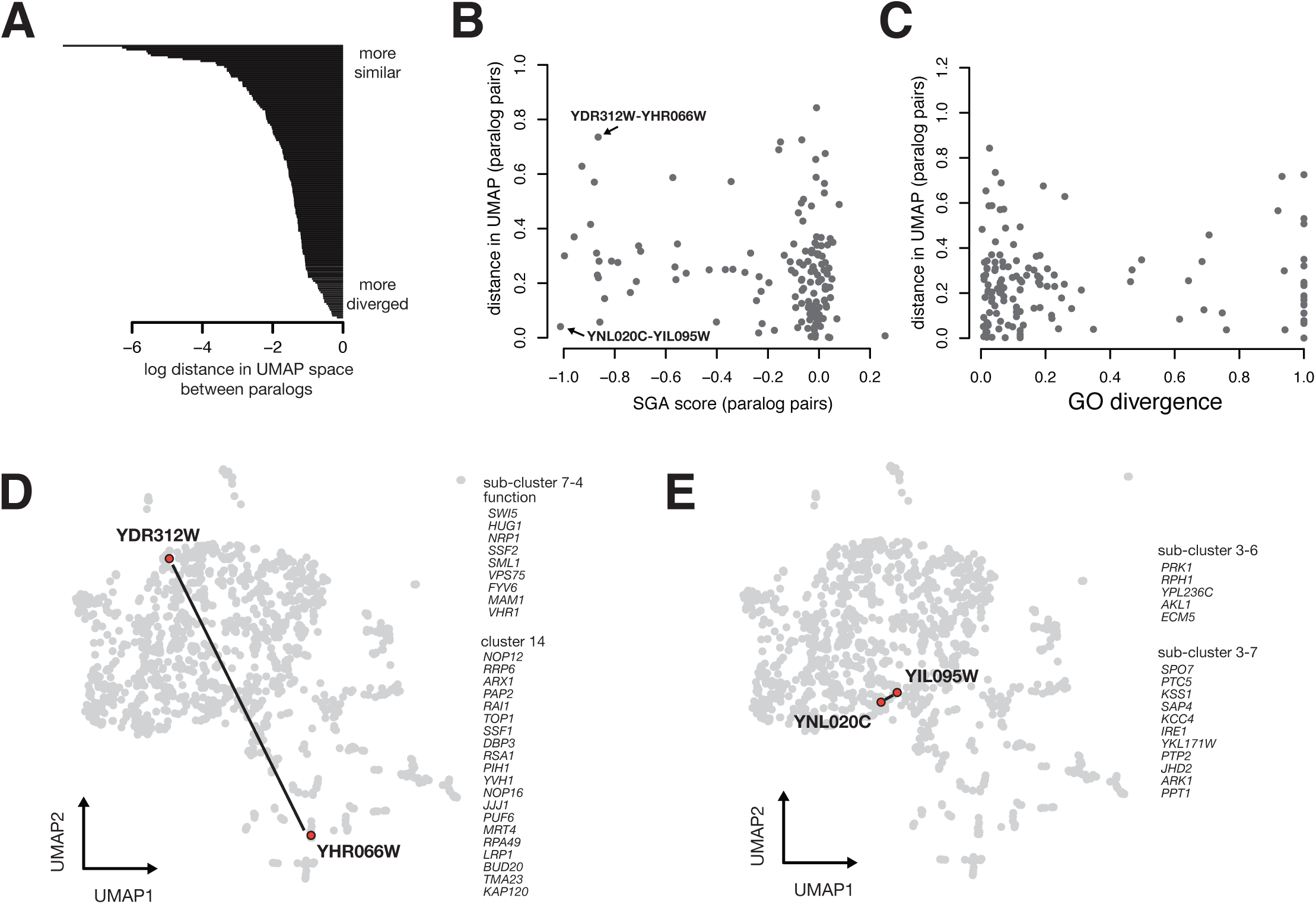
Convergent and divergent function of paralogous gene pairs defined by UMAP distance. (A) Barplot showing log distance in UMAP space between 151 pairs of paralagous gene deletions. (B) Each paralog pair’s UMAP distance plotted against the experimentally-determined synthetic genetic interaction score (briefly, a more negative score on the SGA axis indicates that the double mutant showed a larger cellular fitness defect than the combined additive effect of each single mutants). Two paralog pairs are indicated, and their distance in UMAP space is displayed in (D) and (E). (C) Each paralog pair’s UMAP distance plotted against a metric for paralog divergence calculated using similarlity of GO term annotation. While a low score in the GO divergence metric suggests that paralog pairs have less diverged functions, many of these pairs are far from each other in UMAP space, suggesting that these paralogs show more divergent function than predicted by the GO metric. (D) Full 1484 gene deletion UMAP as in Figure 1A, with a divergent paralog pair (*SSF1* and *SSF2*) highlighted. Genes contained in the same cluster as each paralog are listed; the *SSF1* cluster (7-4) contains many genes required for ribosome biogenesis, while the *SSF2* cluster (14) contains genes involved in DNA damage. (E) Full UMAP with a convergent paralog pair (*ARK1* and *PRK1*) highlighted. Genes contained in the same cluster as each paralog are listed.

